# Auditory cortical responses to speech are shaped by statistical learning of short-term speech input regularities

**DOI:** 10.1101/2022.12.19.520832

**Authors:** Yunan Charles Wu, Vibha Viswanathan, Taylor J. Abel, Lori L. Holt

## Abstract

Speech perception presents an exemplary model of how neurobiological systems flexibly adjust when input departs from the norm. Dialects, accents, and even head colds can negatively impact comprehension by shifting speech from listeners’ expectations. Comprehension improves with exposure to shifted speech regularities, but there is no neurobiological model of this rapid learning. We used electroencephalography to examine human auditory cortical responses to utterances that varied only in fundamental frequency (F0, perceived as voice pitch) as we manipulated the statistical distributions of speech acoustics across listening contexts. Participants overtly categorized speech sampled across two acoustic dimensions that signal /b/ from /p/ (voice onset time [VOT] and F0) to model typical English speech regularities or an expectation-violating accent. These blocks were interleaved with passive exposure to two F0-distinguished test stimuli presented in an oddball ratio to elicit a cortical mismatch negativity (MMN) response. F0 robustly influenced speech categorization when short-term regularities aligned with English but F0 exerted no influence in the context of the accent. The short-term regularities modulated event-related potentials evoked by F0-distinguished test stimuli across both N1 and P3 temporal windows and, for P3 amplitude, there was a strong correlation with perceptual down-weighting of F0. The influence of the short-term regularities persisted to impact MMN in interleaved passive listening blocks when regularities mirrored English but were absent when regularities conveyed the accent. Thus, cortical response is modulated as a function of statistical regularities of the listening context, likely reflecting both early dimension encoding and later categorization.

**Significance Statement:** Speech perception is a quintessential example of how neurobiological systems flexibly adjust when input departs from the norm. Perception is well-tuned to native-language speech patterns. Yet it adjusts when speech diverges from expectations, as with a foreign accent. We observe that the effectiveness of specific cues in speech, like the pitch of a voice, in signaling phonemes like /b/ versus /p/ is dynamically re-weighted when speech violates native-language expectations. We find that this re-weighting is mirrored in cortical responses that reflect both early acoustic dimension encoding and also in later responses linked to phoneme categorization. The results implicate dynamic adjustments in the mapping of speech to cortical representations, as modulated by statistical regularities experienced across local speech input.

## Introduction

Behavior depends upon building representations that respect long-term regularities of the environment. But successful neural systems rapidly adapt to short-term idiosyncratic input that departs from these norms, balancing adaptation to systematic local input statistics without overriding the benefits of alignment with long-term norms.

Speech is a quintessential example of the interplay of stability and adaptive plasticity because its variable acoustics map to representations learned from the long-term speech regularities of a native-language community. Yet, systematic short-term deviations from these norms are as commonplace as chatting with a talker with an unexpected accent or recognizing a spouse’s voice despite their head cold. There is substantial evidence that speech comprehension initially suffers when speech input shifts away from long-term norms, but comprehension improves with exposure to expectation-violating input like a foreign accent (Bertelson et al., 2003; Bradlow & Bent, 2008; Clarke & Garrett, 2004; Greenspan et al., 1988; Hervais-Adelman et al., 2011; Idemaru & Holt, 2011; Maye et al., 2008; Norris et al., 2000; Samuel & Kraljic, 2009; Schwab et al., 1985; Vroomen et al., 2007). Yet, most of what we know has come from behavioral paradigms for which distinct levels of auditory and decisional processes are unavoidably intertwined and impossible to dissociate. There is – as yet – no detailed neurobiological model of how speech perception rapidly adapts to short-term input regularities, like those of an accent (see Ullas et al., 2022 for review).

Dimension-based statistical learning is an attractive paradigm with which to address this gap because it provides a behavioral index of perceptual *weight* – the effectiveness of a specific acoustic dimension in influencing speech categorization. For example, in the context of speech that respects English norms, listeners rely on the acoustic dimension fundamental frequency (F0, experienced as voice pitch) to signal whether a word is *beer* or *pier*; lower frequency F0s better signal *beer* and higher frequency F0s better signal *pier*. But, introduction of an accent that departs from these English norms leads to a rapid shift in how F0 influences speech categorization, such that it is no longer reliably conveys category identity (Idemaru & Holt, 2011, 2014, 2020; Jasmin et al., 2022; Lehet & Holt, 2017, 2020; Liu & Holt, 2015; Schertz et al., 2016; Schertz & Clare, 2019; Wu & Holt, 2022; H. Zhang et al., 2022; X. Zhang & Holt, 2018). Thus, the effectiveness of an acoustic dimension in speech categorization, its *perceptual weight*, dynamically adjusts according to short-term experience with speech regularities that depart from the norm. The mapping from acoustics to speech categories is flexible, not fixed.

Here, we pair dimension-based statistical learning with electroencephalography (EEG) to disentangle processes that may contribute to balancing stability and rapid adaptive plasticity in speech perception. Event-related potentials (ERPs) recorded with EEG during active speech categorization present the opportunity to examine the outcomes of distinct levels of auditory and decisional processing that are intertwined in behavioral response by examining early (N1) and later (P3) auditory evoked potentials that have been linked to graded and more categorical representations of speech, respectively (Pereira et al., 2018; Toscano et al., 2010). We interleave short, active blocks of speech categorization (measuring N1 and P3) in the context of distinct short-term speech input statistics with passive listening blocks. In these passive blocks we leverage the mismatch negativity (MMN) as a marker of the acuity with which acoustic changes are represented by the auditory system (Kujala & Näätänen, 2010). This allows us to ask whether behavioral down-weighting of F0 evident in active blocks carries over to influence cortical sensitivity to acoustic F0 differences in passive listening.

## Materials and Methods

### Participants

Twenty-nine Carnegie Mellon University students (19-30 years; 12 male, 17 female) participated for course credit or pay. All reported normal hearing and American English as the primary language used at home before age two. Equipment malfunction led six participants’ electroencephalography (EEG) data to be unusable. Therefore, behavioral data include 29 participants and EEG analyses include 23 participants. Subjects provided informed consent in accordance with protocols established at Carnegie Mellon University.

### Approach

Figure 1 provides a schematic of the stimuli and approach. Participants first overtly categorized speech stimuli sampled orthogonally from a two-dimensional acoustic space varying in voice onset time (VOT) and fundamental frequency (F0). This provided a baseline behavioral estimate of the perceptual weight with which each acoustic dimension contributes to *beer* versus *pier* categorization. This block proceeded as experimenters placed the electroencephalography (EEG) caps. The remainder of the experiment involved interleaved blocks of overt speech categorization paired with stimulus-evoked EEG (60 trials/block) and passive listening while watching silent film clips during which two speech stimuli with an ambiguous VOT (15 ms) and distinct F0 (220, 300 Hz) were presented in an 85:15 ratio (counterbalanced in assignment across blocks) to elicit a mismatch negativity (MMN) response in the auditory cortical evoked potential (20 trials/block).

**Figure 1.**
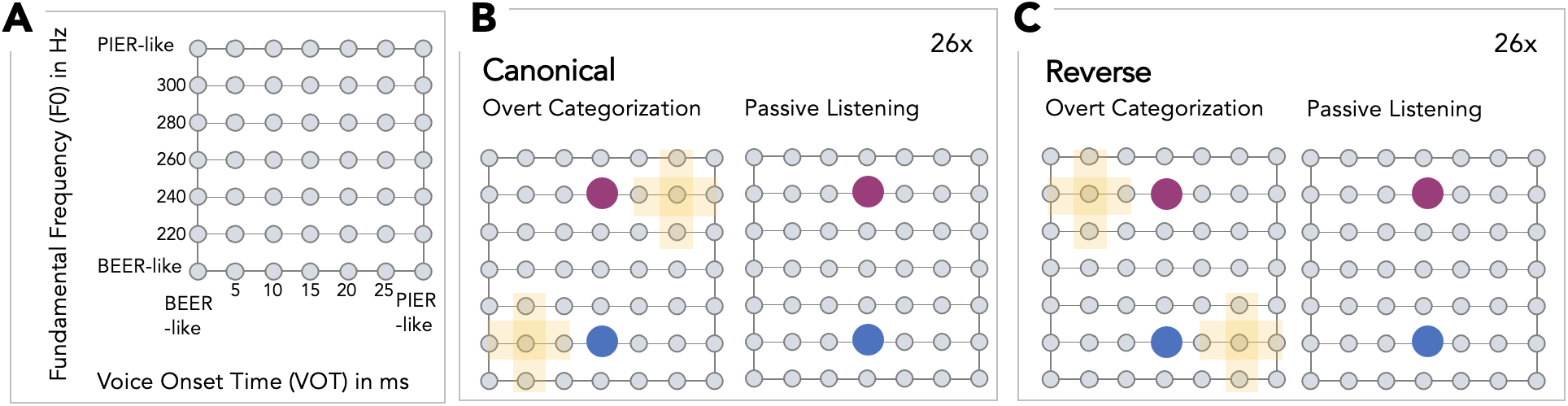
Schematic of stimuli and approach. **A**. Speech stimuli varied acoustically across voice onset time (VOT) and fundamental frequency (F0) and perceptually from *beer* to *pier*, after Idemaru & Holt (2011). Short-term regularities across the F0xVOT acoustic space were created by sampling selectively. **B**. In a block of overt categorization trials with EEG, ten exposure stimuli (yellow) conveyed the Canonical F0xVOT covariation of American English. Test stimuli (magenta, blue) held VOT perceptually ambiguous (15 ms), and varied only in F0 (220, 300 Hz) to assay reliance on F0 in *beer / pier* categorization. This block alternated 26 times with passive listening to only the F0-differentiated test stimuli (magenta, blue), presented in an 85:15 ratio to elicit a mismatch negativity (MMN) response as participants watched silent film clips. Assignment of the two test stimuli as standard (.85 probability) and deviant (.15 probability) was counterbalanced across the blocks. **C**. An ‘accent’ was conveyed by reversing the F0xVOT regularity typical of English in 26 Reverse blocks with passive listening MMN blocks interleaved as described in **B**, and identical F0-differentiated test stimuli.

### Stimuli

The stimuli modeled those of Idemaru and Holt (2011) to create a two-dimensional acoustic space orthogonally sampling voice onset time (VOT, in ms) and fundamental frequency (F0, in Hz), with 7 steps along each acoustic dimension resulting in 49 unique stimuli (Figure 1a).

As in Idemaru and Holt (2011), we used Praat (Boersma, 2006) to manipulate natural productions of *beer* and *pier* spoken by an adult, female monolingual native-English speaker to create 5-ms steps between 0 ms and 30 ms VOT that varied perceptually from *beer* to *pier* (Abramson & Lisker, 1985; Abramson & Whalen, 2017). Fundamental frequency (F0) at vowel onset was linearly interpolated in 20-Hz steps between 200 Hz and 320 Hz to create 7 steps across F0. The F0 contour decreased quadratically to 150 Hz at stimulus offset. Speech stimuli were normalized to the same root mean-squared amplitude.

### Stimulus Presentation and EEG/Behavior Recording

A desktop computer controlled all aspects of the experiment, including triggering sound delivery and storing data. Stimuli (44.1 kHz sampling rate) were presented at 75 dB sound pressure level (SPL) by an RME UFX+ Audio Interface (RME, Haimhausen, Germany) via Etymotic ER-1 (Etymotic, Elk Grove Village, IL) in-ear headphones. The RME sent triggers routed through a S/PDIF-to-TTL converter (Electronics Designs Facility, Boston University) for compatibility with the BioSemi EEG system (Amsterdam, The Netherlands).

The entire experiment was conducted in a double-walled sound-attenuated and electrically shielded booth. A BioSemi ActiveTwo system recorded EEG signals continuously (1024 Hz sampling rate) via Ag/AgCl sintered electrodes embedded in a 32-channel Biosemi Headcap (10-20 system) and left/right mastoid (M1, M2) electrodes. A pair of electrodes placed on the outer canthus of each eye allowed for calculation of the horizontal electrooculogram (EOG) and an additional electrode placed on the left cheek bone allowed for detection of vertical EOG.

An experimenter encouraged participants to remain as still as possible during the experiment to minimize movement artifacts, except to report categorization decisions with a mouse click during overt categorization blocks. Responses were cued by visually presented words (*beer*, pier) on the screen. Messages on the screen encouraged participants to take breaks between blocks.

### Overt Baseline Categorization

The purpose of the baseline categorization block was to assess the perceptual weight of VOT and F0 when short-term input sampled the acoustic space orthogonally, providing no specific F0xVOT covariation. Participants reported whether they heard *beer* or *pier* with a mouse click after the playback of the entire audio file in a baseline block consisting of 25 stimuli (5-25 ms VOT, 220-300 Hz F0; Figure 1a) presented five times each, in a random order (125 total trials). There was no feedback and the ∼5-min task was completed as experimenters placed EEG electrodes.

### Overt Categorization with Canonical/Reverse Short-term Regularities

The purpose of overt speech categorization blocks was to convey short-term speech regularities via Exposure stimuli that align with typical American English F0xVOT patterns (Canonical blocks, Figure 1b) or deviate from typical English patterns (Reverse blocks, Figure 1c), as in an accent. Canonical and Reverse blocks were made up of 10 Exposure stimuli (yellow in Figure 1b-c) that could be unambiguously categorized as *beer* and *pier* via the salient and perceptually unambiguous VOT dimension (Wu & Holt, 2022) and two Test stimuli (Figure 1b) that held VOT perceptually ambiguous (15 ms) with varying F0 (220, 300 Hz). Test stimuli established a scenario in which only F0 was available to signal *beer* versus *pier* and, thus, categorization responses provide a behavioral index of the effectiveness of F0 in directing /b/-/p/ categorization (Idemaru & Holt, 2011; 2014; 2020), its perceptual weight. Each block involved 5 repetitions of each Exposure and Test stimulus for 60 trials paced according to participant responses. Behavioral categorization responses and cortical stimulus-evoked event related potentials measured the impact of the Canonical versus Reverse statistical regularities conveyed by Exposure trials on Test trials that were identical across contexts.

Each participant experienced 26 Canonical blocks interleaved with 30-sec passive listening blocks (see below) and 26 Reverse blocks interleaved with identical passive listening blocks. Canonical and Reverse block order was counterbalanced across participants.

### Passive Listening

The purpose of the passive listening blocks was to examine how the 80-Hz F0 difference across Test stimuli was reflected in the cortical response as a function of the short-term input regularities (Canonical, Reverse) conveyed by interleaved overt speech categorization blocks. Since MMN elicitation requires its own short-term input regularity (an ‘oddball’ paradigm with many Standard stimuli and few Deviants), interleaving overt-categorization and passive-listening blocks allowed us to accumulate an adequate signal-to-noise ratio for measuring MMN without entirely overwriting a potential impact of the (Canonical, Reverse) stimulus statistics that accumulate across overt categorization blocks.

A 20-trial passive listening block immediately followed each 60-trial Canonical or Reverse overt categorization block. In these blocks, participants heard sequences of the two F0-differentiated Test stimuli while watching a silent film with no behavioral task related to the sounds. Participants were instructed to pay attention to the film and ignore the sounds.

Each passive listening block lasted about 30 seconds, with one Test stimulus (counterbalanced across blocks) presented across 17 of 20 trials (.85 probability, Standard) and the other presented 3 times (.15 probability, Deviant). Stimulus order was pseudo-random whereby at least three Standard stimuli preceded a Deviant stimulus. Inter-stimulus interval (ISI) was fixed at 700 ms and the acoustic stream began and ended with 1s of silence.

### EEG Preprocessing

The MNE-PYTHON (Gramfort et al., 2014) open-source software package supported EEG preprocessing. We first down-sampled EEG recordings from each participant to 128 Hz to reduce computational time, then band-pass filtered the result between 0.1 Hz and 32 Hz with a zero-phase finite impulse response (FIR) filter. Next, we performed independent component analysis (ICA), visually inspecting components to remove those generated by blinks, saccades, heartbeats, and muscle movements (Makeig et al., 1996).

Following Toscano et al. (2010) we used the average mastoid (left, right) response to re-reference the preprocessed EEG data for the overt categorization blocks. Next, we epoched the EEG data from 200 ms before stimulus onset to 800 ms after onset and performed baseline correction. We used the average response from frontal electrodes (F3, F4, Fz) to quantify the N1 and the averaged response from parietal electrodes (P3, P4, Pz) to quantify the P3, following prior literature (Toscano et al., 2010; Viswanathan et al., 2021).

To examine the MMN response, we re-referenced the preprocessed data to the average of all channels, then epoched the data from −0.2 s to 0.5 s relative to stimulus onset and performed baseline correction. We used the average response of the frontocentral cluster (electrodes Fz, F3, F4, FC1, FC2, Cz, C3, and C4) to quantify MMN (Moberly et al., 2014; Näätänen et al., 2007a).

### Experimental Design and Statistical Analysis

All participants experienced both Canonical and Reverse speech regularities, allowing us to examine putative behavioral and neural changes within-participants. Statistical analyses focused on behavioral and neural responses to test stimuli, which were constant across listening contexts.

A linear multiple regression model with each participant’s regression weights normalized to sum to one provided an index of baseline behavioral perceptual weight (Holt & Lotto, 2006; Liu & Holt, 2015; Wu & Holt, 2022). We used generalized logistic mixed-effects regression (GLMER) to fit the categorical (*beer* or *pier*) responses (Jaeger, 2008) to (High, Low F0) test stimuli to examine the impact of short-term (Canonical, Reverse) regularities on categorization, with Bayes factor analysis providing a test of the strength of evidence for the predicted interaction across the short-term regularity and test stimulus categorization, a nonparametric independent-samples Mann-Whitney U test to assess the distribution of perceptual weights across short-term regularities, and a one-sample Wilcoxon sign-rank test to examine whether the F0-differentiated test stimuli provide sufficient information to differentiate speech categories.

We performed nonparametric permutation tests (Nichols & Holmes, 2002) to test the null hypotheses that the mean N1 (over the 75-125 ms time window), mean P3 (over the 300-800 ms time window), and MMN (peak Deviant response – peak Standard response; over the 150-350 ms time window) ERPs were the same across Canonical and Reverse contexts. To generate a single sample from each null distribution, we randomly swapped the labels ‘Canonical’ and ‘Reverse’ for the ERP from each participant and computed the average (over participants) ERP difference between Canonical and Reverse contexts. We generated full null distributions for the average n1, P3 and MMN ERP differences between Canonical and Reverse listening contexts by repeating this procedure across 10,000 distinct randomizations. We used these null distributions to assign (one-sided) p-values to the observed average ERP differences between Canonical and Reverse contexts obtained with the correct labels.

We tested the null hypothesis that the MMN wave (Deviant-Standard response time course) is the same across Canonical and Reverse contexts using cluster-based permutation testing (Maris & Oostenveld, 2007) across the 150-350 ms window. We first generated a single sample from the null distribution with ‘Canonical’ and ‘Reverse’ labels randomly swapped for each participant. We then computed the average (over participants) wave across Canonical and Reverse contexts and extracted clusters from the difference wave. We used the null distribution of cluster statistics for the average MMN difference wave (Canonical – Reverse) to assign (one-sided) p-values to the clusters observed in the average MMN difference wave obtained with the correctly labeled data, with a family-wise error rate of .05 as the metric of a significant MMN.

### Code Accessibility

Materials are available at https://osf.io/ysnf3/.

## Results

### Native-English listeners rely on voice onset time (VOT) more than fundamental frequency (F0) in categorizing /b/ versus /p/ at baseline

Figure 2a shows average perceptual weights of VOT and F0 acoustic dimensions in the baseline block, calculated as predictor variables in a linear multiple regression model and normalized such that each participant’s regression weights to sum to one (Holt & Lotto, 2006). These patterns confirm expectations that native American-English listeners rely most on VOT in /b/-/p/ categorization although F0 plays a secondary role in signaling category identity (Holt et al., 2018; Idemaru & Holt, 2011; Winn et al., 2013; Wu & Holt, 2022).

**Figure 2.**
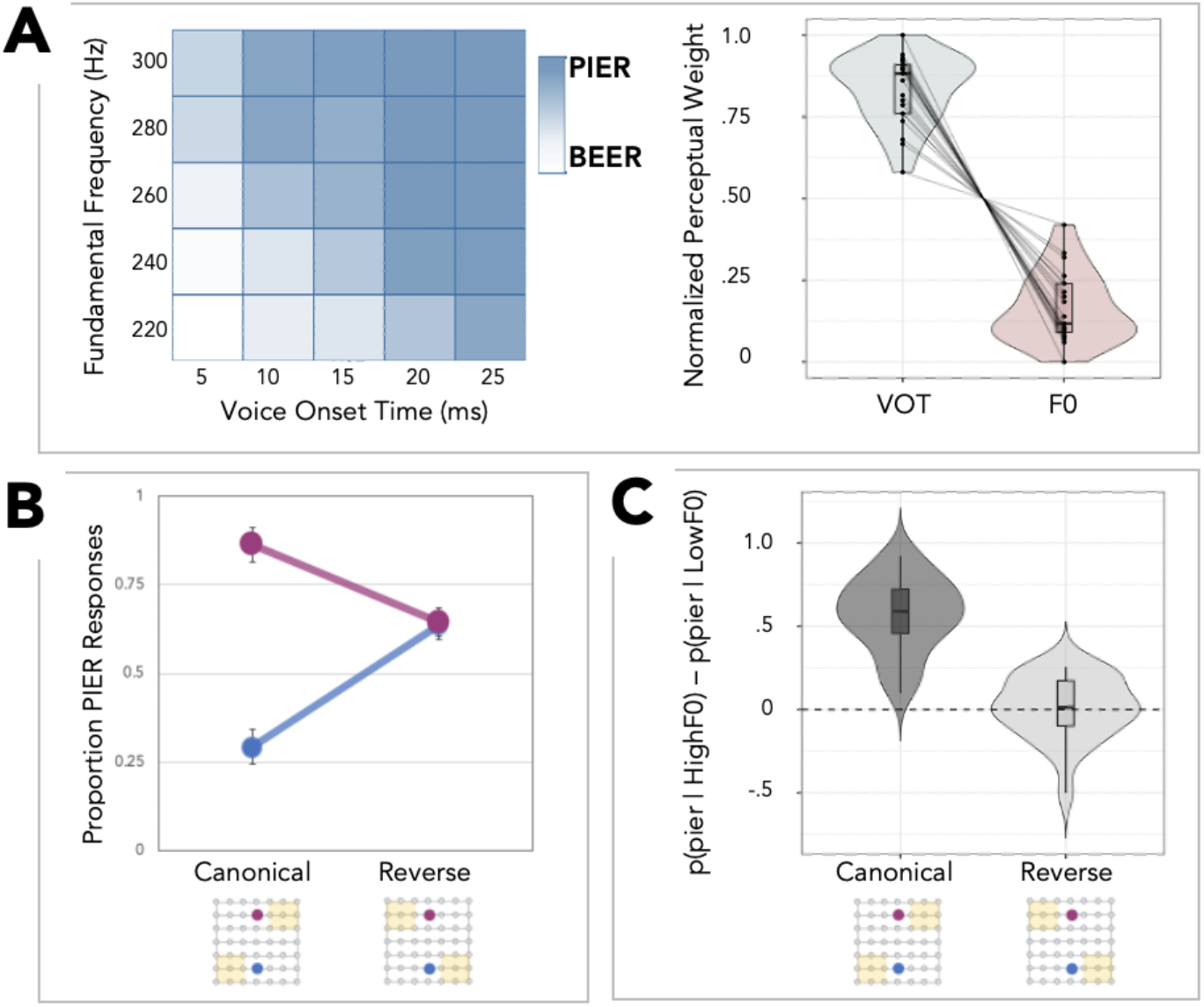
Overt Categorization in the Baseline and Canonical/Reverse Blocks. **A**. At baseline, listeners used VOT as the primary dimension in categorizing *beer* and *pier*, with F0 playing a secondary role. The heatmap presents average Baseline *beer* (white) versus *pier* (blue) categorization responses across an acoustic space orthogonally sampling F0 and VOT. The same data are shown as combined box and violin plots (outliers trimmed in the violin plots) as normalized dimension weights for VOT (left) and F0 (right); lines indicate individual participants. **B**. In the context of short-term regularities that violate expectations, the influence of F0 in categorization is down-weighted. Average percent *pier* responses to the F0-differentiated test stimuli in Canonical and Reverse blocks; error bars indicate standard errors. Magenta indicates responses for to the High F0 test stimulus and blue indicates responses to the Low F0 test stimulus. **C**. The distributions of response differences across test stimuli (p(*pier*|High F0) -p(*pier*|Low F0)) for the Canonical and Reverse blocks illustrate that listeners rely on F0 less for *beer-pier* categorization in the context of Reverse short-term speech input regularities. In both panels A and C, the median (horizontal bar), 50% confidence limits (black box), and 95% confidence limits (black whiskers) are indicated for each box plot.

### Short-term speech input regularities shift baseline perceptual weights

Figure 2b shows the average proportion *pier* responses of F0-differentiated test stimuli across Canonical and Reverse blocks. Using generalized logistic mixed-effects regression (GLMER) to fit the categorical (*beer* or *pier*) responses (Jaeger, 2008), we constructed two GLMER models with participants as a random effect: a base model with the main effect of F0-differentiated test stimulus and the main effect of short-term regularity as a function of Block (AIC = 8089.2, BIC = 8116.7), and a full model with each main effect and their interaction. The two models indicated a significant interaction between F0 and Block (likelihood ratio test, with chi-square approximation: p < 0.001, AIC = 7361.0, BIC = 7395.3, χ^2^ = 730.27). A Bayes factor analysis also revealed strong evidence for the full model versus the base model (Bayes Factor > 100). Introduction of the accent in the Reverse block resulted in down-weighting of the influence of F0 in categorization, consistent with prior research (Idemaru & Holt, 2011, 2014, 2020; Lehet & Holt, 2017, 2020; Liu & Holt, 2015; X. Zhang et al., 2021; X. Zhang & Holt, 2018).

Figure 2c shows the significantly different distributions of F0 perceptual weights across Canonical and Reverse blocks (nonparametric independent-samples Mann-Whitney U test, W = 514, p < 0.001). A one-sample Wilcoxon sign-rank test established that F0 provides sufficient information differentiate *beer* from *pier* in Canonical blocks ([p(*pier*|High F0) minus p(*pier*)|Low F0] > 0, V = 276, p < 0.001) but not in Reverse blocks (V = 145, p = 0.28).

### The MMN response to acoustically distinct speech stimuli is eliminated when short-term regularities violate English norms

The MMN response, characterized by significantly stronger negativity for a deviant, compared to a standard stimulus, is thought to signal acoustic change detection or discriminability (Näätänen et al., 2007) and can inform levels of cortical processing with its time course (Moberly et al., 2014). Drawing from a rich MMN literature (Fitzgerald & Todd, 2020) including a study of the perceptual weight of acoustic dimensions conveying speech categories (Moberly et al., 2014), we predicted that MMN amplitude would be modulated by short-term speech regularities experienced in interleaved overt categorization blocks. We specifically expected that F0-differentiated test stimuli would elicit larger MMN magnitudes in the context of interleaved Canonical blocks compared to interleaved Reverse blocks.

Figure 3a shows the average stimulus-evoked EEG response from the frontocentral cluster (Fz, F3, F4, FC1, FC2, Cz, C3, and C4) (Moberly et al., 2014; Näätänen et al., 2007) for standard versus deviant F0-differentiated test stimuli in the context of interleaved Canonical versus Reverse blocks conveying distinct short-term speech regularities. Figure 3b shows the MMN (difference of deviant compared to standard, with each F0-differentiated test stimuli serving in each role), across contexts.

**Figure 3.**
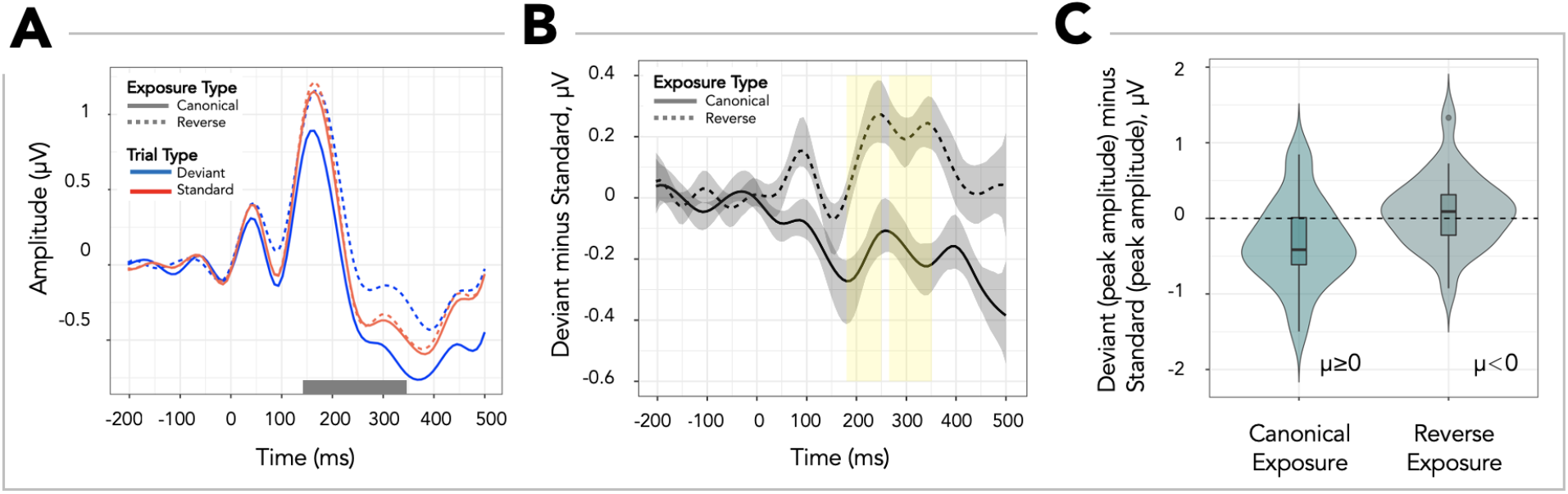
MMN response. Event-related potentials (ERPs) time-locked to onset of the High and Low F0 test stimuli presented in the passive listening blocks, as a function of whether interleaved blocks with overt categorization decisions had Canonical or Reverse short-term regularities. **A**. Standard and Deviant event-related potentials (ERPs, red versus blue lines, respectively) from the Canonical and Reverse contexts (solid versus dashed lines). The grey shaded region on the x-axis indicates the 150-350 ms temporal window across which peak amplitudes in Standard and Deviant ERPs were extracted, following Moberly et al. (2014). **B**. Smoothed MMN response waves (amplitude difference, Deviant - Standard) for the Canonical and Reverse blocks. Grey shaded areas indicate bootstrapped 95% confidence bands (Shalizi, 2013; nonparametric quantile bootstrap across participants) around the ERP; curves were smoothed using spline smoothing with 15 knots. Yellow shaded regions indicate time points for which the Canonical MMN response wave (Deviant - Standard) was significantly different from the MMN elicited by the same stimuli in the Reverse context. **C**. Distribution of the difference in ERP peak amplitude between test stimuli presented as low-probability Deviants and those presented as high-probability Standards following exposure to Canonical, and separately, Reverse short-term speech input regularities (combined box and violin plots with outliers trimmed in the violin plots). The median (horizontal bar), 50% confidence limits (black box), and 95% confidence limits (black whiskers) are indicated for each box plot. The dashed line indicates no difference, i.e., no MMN response.

Moberly et al. (2014) reported an early temporal window (201-250 ms) across which MMN reflects sensory dimensions of speech input and a later window (251-300 ms) reflecting the perceptual weight associated with the effectiveness of dimensions in signaling speech categories. Accordingly, here we examined MMN in the 150-350 ms time window as a function of short-term speech input regularity by performing a permutation test on the MMN wave (Deviant-Standard response time course) comparing Canonical and Reverse contexts. Figure 3b illustrates time clusters with significant differences (nonparametric cluster-based permutation test with alpha threshold .05; Maris & Oostenveld, 2007) in MMN across Canonical and Reverse contexts; significant MMN response differences were found in two time-clusters: 180-250 ms, and 266-352 ms, which are wider than, but approximately coincide with, the early MMN temporal windows observed by Moberly et al. (2014).

Figure 3c shows the across-participant distribution of the difference between the deviant and standard peak amplitudes, with each peak amplitude computed over the 150-350 ms time window. The dashed line in the figure indicates no MMN. A nonparametric permutation test revealed a marginally stronger negative peak for deviant than for standard in the context of interleaved Canonical blocks (Deviant-Standard: μ < 0, p = 0.055) that was not observed when interleaved blocks conveyed the Reverse regularity (Deviant-Standard: μ ≥ 0, p = 0.70).

The MMN response, typically understood to be a pre-attentive response to acoustically distinct input (Näätänen & Alho, 1997), is modulated according to short-term input regularities in recently experienced speech. Two speech stimuli differentiated by a substantial 80-Hz F0 difference elicit a significant MMN when interleaved regularities align with expectations from English, but do not when regularities violate English norms.

### N1 amplitude differences across acoustically distinct speech stimuli are reduced when local speech regularities depart from the norm

The N1 is a large, negative potential distributed mostly over the frontocentral scalp electrodes about 80 and 120 milliseconds after stimulus onset (Näätänen & Picton, 1987). Its amplitude is modulated by within-category acoustic differences of speech sounds, positioning it as a general metric of perceptual encoding at early stages of perception across a wide range of speech contrasts (Pereira et al., 2018; Toscano et al., 2010) We calculated the mean ERPs for the N1 across frontal electrodes (F3, F4, Fz) separately for Canonical and Reverse blocks in the 75-125 ms window (Toscano et al., 2010) and fit a linear mixed-effects model (LMER) to the N1 peak amplitude.

Figure 4a shows that the across-participants distributions of the mean amplitude difference for the N1 response elicited by the F0-differentiated test stimuli were significantly different across Canonical and Reverse short-term regularities (nonparametric permutation test; p = .02).

**Figure 4.**
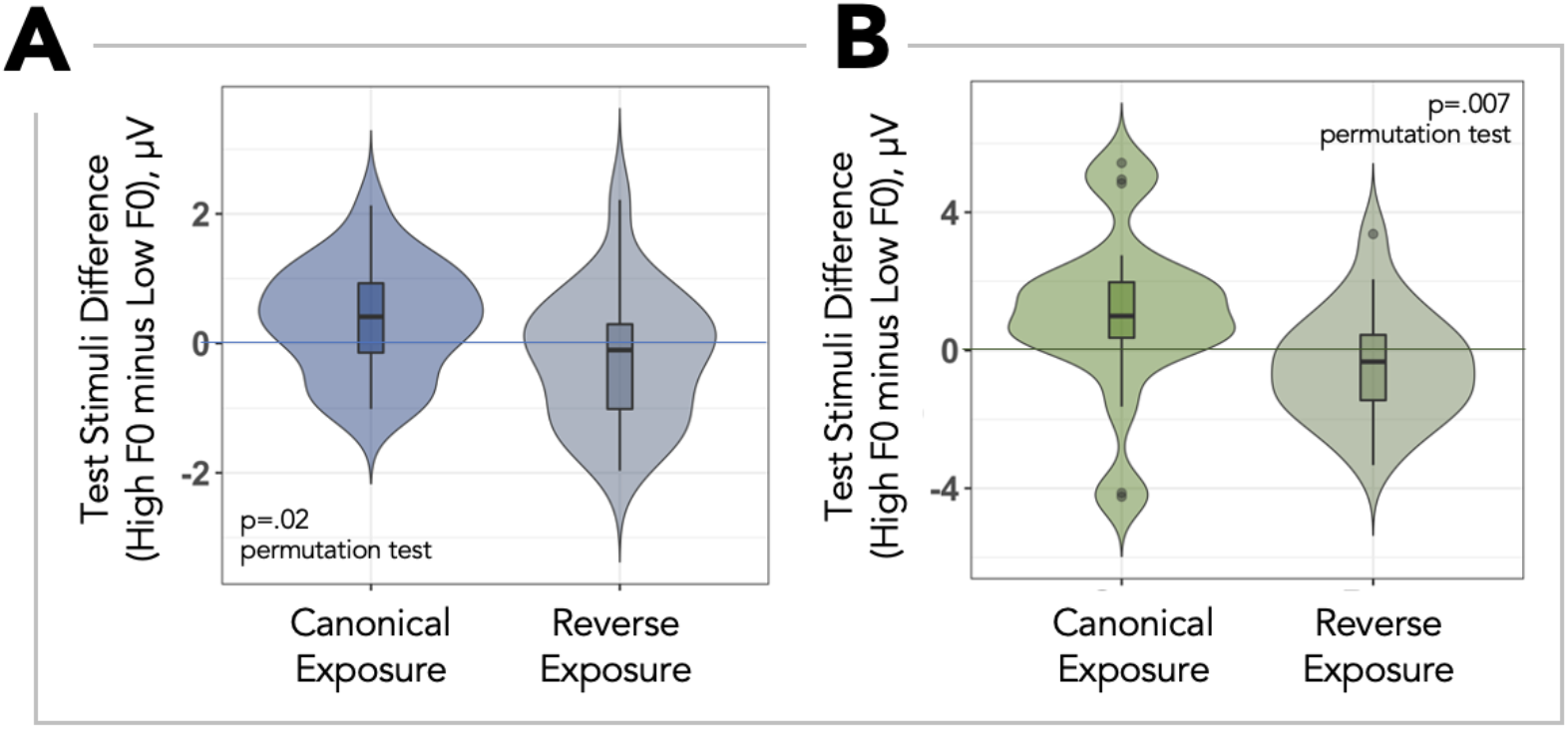
ERP analysis in overt categorization with Canonical versus Reverse regularities. **A**. Distribution of difference (High F0 - Low F0) in N1 response (averaged across a 75-125 ms temporal window) for the test stimuli differentiated by F0 for the Canonical and Reverse blocks (combined box and violin plots with outliers trimmed in the violin plots). **B**. Distribution of the difference (High F0 - Low F0) in P3 response (averaged across a 300-800 ms temporal window) for the test stimuli differentiated by F0 for the Canonical and Reverse blocks (combined box and violin plots with outliers trimmed in the violin plots). P-values for the permutation tests comparing Canonical and Reverse contexts are also indicated. In both panels A and B, the median (horizontal bar), 50% confidence limits (black box), and 95% confidence limits (black whiskers) are indicated for each box plot.

Examining N1 response amplitude with a base model included main effects of the F0-differentiated test stimuli, the short-term regularities conveyed across blocks, and participant (random effect) as predictors (AIC = 316.18, BIC = 328.78). Comparing this base model with a full model including the interaction revealed a significant interaction between block (Canonical, Reverse) and test stimulus F0 (High, Low) (Likelihood ratio test, with chi-square approximation: p = 0.034, AIC = 313.17, BIC = 328.30, chi-square = 5.01). Moreover, Bayes factor analysis of the two models provided substantial evidence for the full model versus the base model (Bayes Factor = 12.22).

### P3 amplitude differences across acoustically distinct speech stimuli are reduced when local speech regularities depart from the norm

The P3 is characteristically associated with stimulus evaluation, categorization or decision making. Evoked by speech, it aligns better with category differences than acoustic change (Dalebout & Stack, 1999; Dehaene-Lambertz, 1997; Horev et al., 2007; Maiste et al., 1995; Toscano et al., 2010). We calculated the mean ERPs for the P3 across parietal electrodes (P3, P4, Pz) and over a 300-800 ms window (Toscano et al. 2010), separately for Canonical and Reverse contexts. Figure 4b shows that the across-participants distributions of the mean amplitude difference for the P3 response elicited by the F0-differentiated test stimuli were significantly different across Canonical and Reverse short-term regularities (nonparametric permutation test; p = .007). We then fit a linear mixed-effects model (LMER) to the mean P3 amplitude with a base model that included main effects of the F0-differentiated test stimuli, the short-term regularities conveyed across blocks, and participant (random effect) as predictors (AIC = 396.72, BIC = 409.33). Comparing this base model with a full model including the interaction revealed a significant interaction between block (Canonical, Reverse) and test stimulus F0 (High, Low) (Likelihood ratio test, with chi-square approximation: p = 0.03, AIC = 393.32, BIC = 408.45, χ^2^= 5.4046). Moreover, Bayes factor analysis of the two models provided substantial evidence for the full model versus the base model (Bayes Factor = 21.80).

### Categorization accuracy in the context of the accent predicts the degree of F0 down-weighting in behavior

Figure 5a illustrates a significant negative relationship between exposure trial categorization accuracy (according to the salient VOT dimension) in the Reverse context and the behavioral weighting of F0 (p(*pier*|LowF0) – p(*pier*|HighF0) as quantified from the behavioral difference in labeling F0-differentiated test trials (r = −.479, p = .02), replicating Wu and Holt (2022). Greater accuracy in labeling exposure trials in the Reverse blocks (and, by hypothesis, greater differentiation in category activation) was associated with greater down-weighting of F0 in speech categorization. The more accurately participants categorized exposure trials conveying the accent, the greater the degree of F0-downweighting. This implicates category activation as a driver of the dynamic behavioral and cortical adjustments in response to speech regularities that violate expectations.

**Figure 5.**
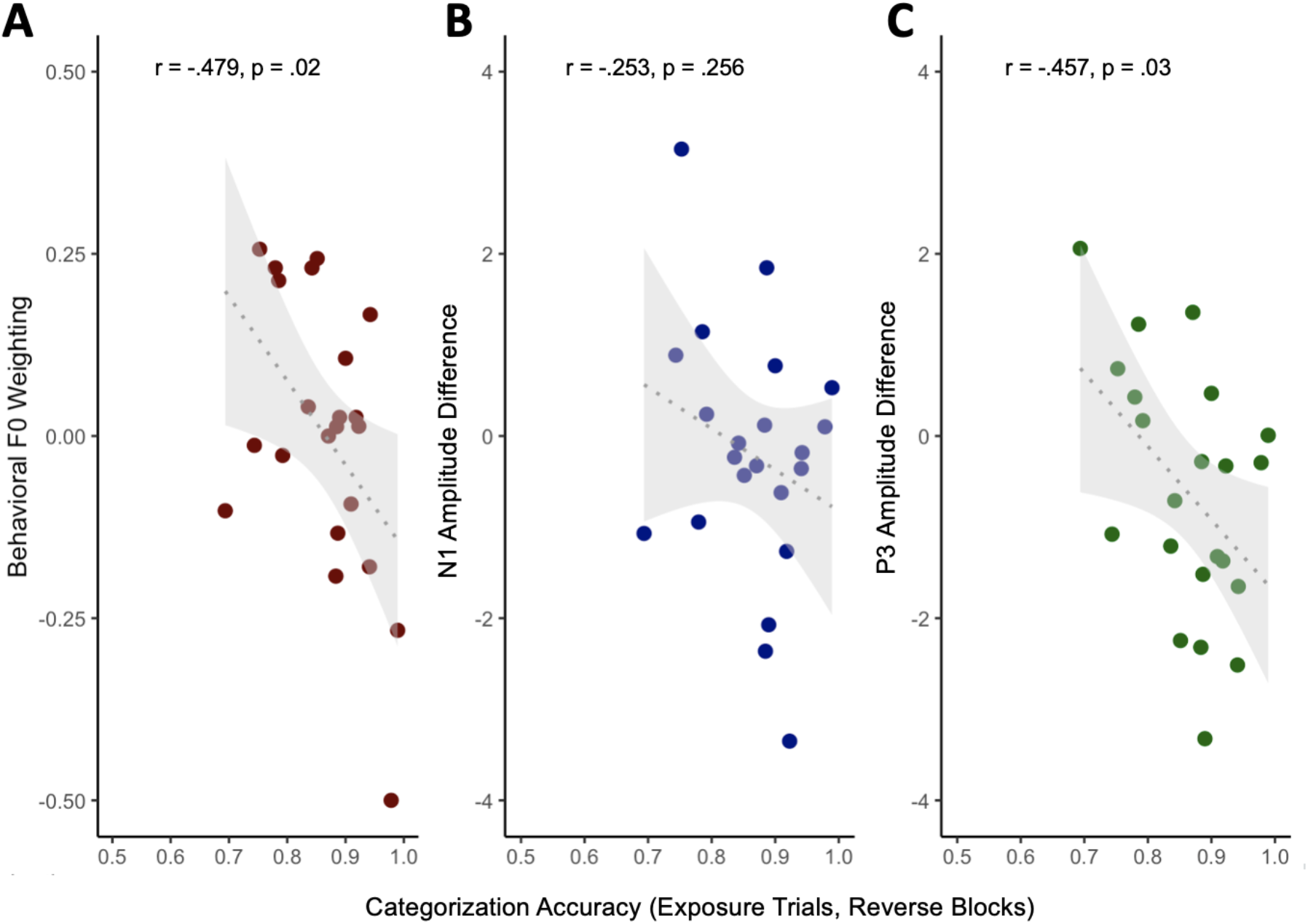
Relationship of categorization accuracy of exposure trials in Reverse blocks to behavioral and event related potential measures. **A**. Exposure accuracy versus the behavioral weighting of F0 (p(pier | HighF0) – p(pier | LowF0)) across Test stimuli in the Reverse blocks. **B**. Exposure accuracy versus mean N1 amplitude evoked by High F0 versus Low F0 Test stimuli. **C**. Exposure accuracy versus mean P3 amplitude evoked by High F0 versus Low F0 Test stimuli. The shaded gray region indicates the 95% confidence interval. Correlations are presented across N=23 participants who contributed both behavioral and EEG data. Across participants, average exposure trial accuracy was 84.76% (SD=10.7%). We rejected one participant because exposure accuracy fell 2 SD from the mean (50.38%). Thus, 22 participants are included here.

### Categorization accuracy in the context of the accent predicts smaller P3 amplitude differences across F0-differentiated test stimuli

Figures 5b and 5c plot the relationship between exposure trial categorization accuracy in the Reverse context and the differences for the N1 and P3 elicited by F0-differentiated test stimuli. Accuracy in categorizing the exposure trials in the Reverse block according to the primary acoustic input dimension (here, VOT) was significantly associated with lower-amplitude P3 responses (r = −.457, p = .03), but there was no relationship with the N1 response (r = −.253, p = .256).

## Discussion

Neural systems must balance stability and plasticity. As an ecologically significant example, speech acoustics map to representations learned from long-term native-language norms, but deviations from these norms are common. We leveraged electroencephalography to ask how the cortical response to speech varies according to the statistics of a listening context.

Our testbed was *beer* versus *pier* speech categorization for which two acoustic dimensions contribute substantially: VOT (related to the length of time from lip release to the onset of voicing from vocal fold vibration) and F0 (the frequency of that vibration). American-English listeners rely most on VOT to inform category decisions and, secondarily, on F0 (Wu & Holt, 2022). Yet, upon encountering an ‘accent’ that reverses the typical English relationship between VOT and F0, F0 no longer reliably signals category identity. This dimension-based statistical learning highlights the perceptual system’s sensitivity to long-term input norms, and its rapid adjustment when input violates these expectations (Idemaru & Holt, 2011; 2014; 2020; Liu & Holt, 2015; Lehet & Holt, 2017; 2020; Zhang & Holt, 2018; Zhang et al., 2021; Schertz et al. 2016; Wu & Holt, 2022; Hodson et al., 2022; Jasmin et al. 2022). It is worth emphasizing that here, as in prior studies, the rapid adjustment of the perceptual weight of F0 in informing speech categorization arose implicitly and in the context of statistical regularities conveyed by a single voice.

The influence of differing statistical regularities in speech persisted to affect subsequent cortical responses in passive listening, as indexed by the mismatch negativity (MMN) response. This is revealing in that MMN amplitude is taken as a marker of the acuity with which acoustic changes are represented by the auditory system (Fitzgerald & Todd, 2020; Garrido et al., 2009; Näätänen, 2008; Näätänen et al., 2007b). Here, this acuity was modulated according to interleaved listening contexts with distinct statistical regularities even though the acuity was measured across a substantial 80-Hz F0 difference. The interleaved statistical regularities of our design impacted the MMN in each of the temporal windows identified by Moberly et al. (2014), consistent with an influence of statistical listening contexts on both perceptual weights of input dimensions (later window) and dimension encoding (earlier window). This is of interest in that there are distinct later (frontal) and earlier (bilateral auditory cortex) cortical generators of MMN (Alho, 1995). At the broadest level, the MMN results indicate that the potentially pre-attentive auditory cortical evoked response measured as the MMN is dynamically and rapidly influenced by short-term experienced in speech, such as in an accent.

The results demonstrate the utility of the novel approach we took to measuring MMN. Elicitation of the MMN response depends upon input regularities that convey a stimulus (or set of stimuli) as ‘standard’ in contrast to a lower probability ‘deviant’. This makes MMN rather unsuitable for examining statistical learning phenomena that are not specifically tied to stimulus probability. Our approach was to interleave brief passive listening blocks with probabilities appropriate to eliciting the MMN with blocks of trials conveying regularities expected to evoke statistical learning. Cycling between blocks conveying the canonical English regularity or the accent and brief MMN-eliciting passive listening blocks proved successful in eliciting the MMN and in modulating it according to the interleaved statistical context. Relatedly, the carryover of influence from active categorization blocks to passive listening in MMN blocks allows us to conclude that the behavioral down-weighting of F0 is consistent with a persistent learning effect rather than mere modulation of activation, and that this learning is robust across listening contexts because it carries over from active categorization to influence the neural response in passive listening.

Short-term statistical speech regularities also modulated ERPs elicited by F0-differentiated test stimuli in the context of active speech categorization. For both N1 and P3, ERP amplitude differences across test stimuli were evident when regularities aligned with English but were ameliorated when regularities violated expectations.

Multiple cortical generators may contribute to the N1 response (Naatanen & Picton, 1987). The dominant contributors to the N1 have been reported to be the bilateral transverse temporal gyri (Heschl’s gyri) with contributions from other regions depending on stimulus and listening condition (Scherg and Von Cramon, 1984; Pantev et al., 1989). The amplitude of the N1 has been argued to be a general index of perceptual encoding at early stages of auditory perception across a range of phonetic contrasts in speech (Toscano, McMurray, Dennhardt, & Luck, 2010; Pereira, Gao, & Toscano, 2018) and thereby may convey a rather veridical representation of input acoustics. In this context, the amelioration of the N1 amplitude difference when statistical regularities run counter to English norms would indicate that dimension-based statistical learning impacts the encoding of F0. The present results are consistent with an understanding of cortical feature encoding as dynamically context-dependent according to statistical regularities experienced across speech input.

Prior studies have associated P3 amplitude with post-perceptual categorization processing whereby P3 amplitude is associated with category differences, rather than acoustic changes (Dalebout & Stack, 1999; Dehaene-Lambertz, 1997; Maiste et al., 1995; Toscano et al., 2010). In general, P3 responses in active speech categorization tasks appear to originate across a network of brain regions involving frontal cortical regions areas including the anterior and posterior portions of inferior frontal gyrus (Toscano et al., 2018). Other studies suggest that this is but one node of a distributed network contributing to P3 including medial temporal lobe, thalamus, superior temporal sulcus, ventrolateral prefrontal cortex, and posterior superior parietal cortex (Halgren et al., 1980, 1998; Mulert et al., 2004; Nieuwenhuis et al., 2005; Yingling & Hosobuchi, 1984).

Our F0-differentiated test stimuli elicited distinct P3 responses, with high F0 test stimuli associated with *pier* responses producing greater-amplitude P3s than low F0 test stimuli in the blocks of trials conveying regularities typical of English speech. Introduction of the accent eliminated these P3 amplitude differences for F0-differentiated test stimuli, mirroring category decisions in behavior. In this sense, modulation of P3 may reflect the adaptive effects of short-term listening context on speech categorization such that it is elicited by pre-categorical influence on the effectiveness of F0 in driving differentiated categorization, as is evident in MMN and N1 modulation.

This interpretation is supported by individual differences in participants’ susceptibility to the short-term regularities conveyed by accent. As observed in prior research (Wu & Holt, 2022), participants who were most successful at categorizing exposure stimuli conveying the accent (according to VOT) exhibited the most robust down-weighting of F0. The relationship of the amplitude difference of the N1 response to the same stimuli with exposure stimulus categorization trended in the same direction but was not significant. In contrast, P3 amplitude differences across F0-differentiated test stimuli – putatively aligned with category-level processing – exhibit this same association. In general, the more successfully participants used the VOT dimension to categorize exposure trials conveying the accent, the greater the magnitude of neural (and behavioral) down-weighting across test stimuli differentiated only by F0.

The robustness of the association of category activation (indexed by exposure trial categorization success in the Reverse blocks) with the magnitude of F0-downweighting is informative to theory. According to an error-driven learning account (Wu & Holt, 2022), successful activation of category representations via unambiguous VOT for Reverse block exposure stimuli may trigger predictions about typical F0 patterns that align with long-term norms. When input mismatches these predictions, as in Reverse blocks, an error signal may facilitate a shift in the encoding or mapping of the secondary, F0, dimension. This may lead to down-weighting of the F0 dimension we observe behaviorally, as well as the response differences in the cortical responses we observe in the current study.

In summary, cortical response is rapidly impacted by short-term speech input statistics. When relationships among acoustic input dimensions violate long-term language norms, as for an accent, the effectiveness of acoustic dimensions in speech categorization is dynamically adjusted. The present results suggest that these adjustments arise from an influence on both the neural encoding of dimensions (which may be reflected in early MMN temporal windows and the N1), and acoustic-phonetic mapping that ultimately impact decision-driven categorization response (which may be reflected in the P3). This is consistent with a model of cortical representation of speech input that is flexible, not fixed.

## Acknowledgements

The work was supported by a doctoral dissertation grant to LLH and YCW from the National Science Foundation (BCS1941357), and by awards from the National Institutes of Health to LLH and TJA (R21DC019217) and to VV (T32DC11499, F32DC020649). We appreciate the engagement of Prof. Barbara Shinn-Cunningham’s laboratory as part of the Pittsburgh Cognitive Auditory Neuroscience collective. Data and code are available at https://osf.io/ysnf3/.

## References

Abramson, A., & Lisker, L. (1985). Relative power of cues: F0 shift versus voice timing.

Abramson, A. S., & Whalen, D. H. (2017). Voice Onset Time (VOT) at 50: Theoretical and practical issues in measuring voicing distinctions. Journal of Phonetics, 63, 75–86. https://doi.org/10.1016/j.wocn.2017.05.002

Alho, K. (1995). Cerebral Generators of Mismatch Negativity (MMN) and Its Magnetic Counterpart (MMNm) Elicited by Sound Changes. Ear and Hearing, 16(1), 38–51. https://doi.org/10.1097/00003446-199502000-00004

Benjamini, Y., & Hochberg, Y. (1995). Controlling the false discovery rate: a practical and powerful approach to multiple testing. Journal of the Royal Statistical Society, Series B (Methodological), 289–300.

Bertelson, P., Vroomen, J., & Gelder, B. (2003). Visual recalibration of auditory speech identification: a McGurk aftereffect. Psychological Science, 14(6), 592–597. https://doi.org/10.1046/j.0956-7976.2003.psci_1470.x

Boersma, P. (2006). Praat: doing phonetics by computer.

Bradlow, A. R., & Bent, T. (2008). Perceptual adaptation to non-native speech. Cognition, 106(2), 707–729. https://doi.org/10.1016/j.cognition.2007.04.005

Clarke, C. M., & Garrett, M. F. (2004). Rapid adaptation to foreign-accented English. The Journal of the Acoustical Society of America, 116(6), 3647–3658. https://doi.org/10.1121/1.1815131

Dalebout, S. D., & Stack, J. W. (1999). Mismatch negativity to acoustic differences not differentiated behaviorally. Journal of the American Academy of Audiology, 10(7), 388–399.

Dehaene-Lambertz, G. (1997). Electrophysiological correlates of categorical phoneme perception in adults. NeuroReport, 8(4), 919–924. https://doi.org/10.1097/00001756-199703030-00021

Fitzgerald, K., & Todd, J. (2020). Making Sense of Mismatch Negativity. Frontiers in Psychiatry, 11, 468. https://doi.org/10.3389/fpsyt.2020.00468

Garrido, M. I., Kilner, J. M., Stephan, K. E., & Friston, K. J. (2009). The mismatch negativity: A review of underlying mechanisms. Clinical Neurophysiology, 120(3), 453–463. https://doi.org/10.1016/j.clinph.2008.11.029

Godey, B., Schwartz, D., Graaf, J. B. de, Chauvel, P., & Liégeois-Chauvel, C. (2001). Neuromagnetic source localization of auditory evoked fields and intracerebral evoked potentials: a comparison of data in the same patients. Clinical Neurophysiology, 112(10), 1850–1859. https://doi.org/10.1016/s1388-2457(01)00636-8

Gramfort, A., Luessi, M., Larson, E., Engemann, D. A., Strohmeier, D., Brodbeck, C., Parkkonen, L., & Hämäläinen, M. S. (2014). MNE software for processing MEG and EEG data. NeuroImage, 86, 446–460. https://doi.org/10.1016/j.neuroimage.2013.10.027

Greenspan, S. L., Nusbaum, H. C., & Pisoni, D. B. (1988). Perceptual Learning of Synthetic Speech Produced by Rule. Journal of Experimental Psychology: Learning, Memory, and Cognition, 14(3), 421–433. https://doi.org/10.1037/0278-7393.14.3.421

Halgren, E., Marinkovic, K., & Chauvel, P. (1998). Generators of the late cognitive potentials in auditory and visual oddball tasks. Electroencephalography and Clinical Neurophysiology, 106(2), 156–164. https://doi.org/10.1016/s0013-4694(97)00119-3

Halgren, E., Squires, N. K., Wilson, C. L., Rohrbaugh, J. W., Babb, T. L., & Crandall, P. H. (1980). Endogenous Potentials Generated in the Human Hippocampal Formation and Amygdala by Infrequent Events. Science, 210(4471), 803–805. https://doi.org/10.1126/science.7434000

Hervais-Adelman, A. G., Davis, M. H., Johnsrude, I. S., Taylor, K. J., & Carlyon, R. P. (2011). Generalization of Perceptual Learning of Vocoded Speech. Journal of Experimental Psychology: Human Perception and Performance, 37(1), 283–295. https://doi.org/10.1037/a0020772

Hodson, A. J., Shinn-Cunningham, B., & Holt, L. L. (2022, November 21). Statistical learning across passive listening adjusts perceptual weights of speech input dimensions. https://doi.org/10.31234/osf.io/4kxz3

Holt, L. L., & Lotto, A. J. (2006). Cue weighting in auditory categorization: implications for first and second language acquisition. The Journal of the Acoustical Society of America, 119(5 Pt 1), 3059–3071. https://doi.org/10.1121/1.2188377

Holt, L. L., Tierney, A. T., Guerra, G., Laffere, A., & Dick, F. (2018). Dimension-selective attention as a possible driver of dynamic, context-dependent re-weighting in speech processing. Hearing Research, 366, 50–64. https://doi.org/10.1016/j.heares.2018.06.014

Horev, N., Most, T., & Pratt, H. (2007). Erratum&colon; “Olivocochlear Efferents&colon; Anatomy, Physiology, Function, and the Measurement of Efferent Effects in Humans” &lsqb;Ear and Hearing, 27(6)&colon;589–607 (2006)&rsqb; Ear and Hearing, 28(1), 129. https://doi.org/10.1097/01.aud.0000250021.69163.96

Idemaru, K., & Holt, L. L. (2011). Word recognition reflects dimension-based statistical learning. Journal of Experimental Psychology: Human Perception and Performance, 37(6), 1939–1956. https://doi.org/10.1037/a0025641

Idemaru, K., & Holt, L. L. (2014). Specificity of dimension-based statistical learning in word recognition. Journal of Experimental Psychology. Human Perception and Performance, 40(3), 1009–1021. https://doi.org/10.1037/a0035269

Idemaru, K., & Holt, L. L. (2020). Generalization of dimension-based statistical learning. Attention, Perception, & Psychophysics, 1–19. https://doi.org/10.3758/s13414-019-01956-5

Jasmin, K., Tierney, A., Obasih, C., & Holt, L. (2022). Short-term perceptual reweighting in suprasegmental categorization. Psychonomic Bulletin & Review, 1–10. https://doi.org/10.3758/s13423-022-02146-5

Kujala, T., & Näätänen, R. (2010). The adaptive brain: A neurophysiological perspective. Progress in Neurobiology, 91(1), 55–67. https://doi.org/10.1016/j.pneurobio.2010.01.006

Lehet, M., & Holt, L. L. (2017). Dimension-Based Statistical Learning Affects Both Speech Perception and Production. Cognitive Science, 41 Suppl 4, 885–912. https://doi.org/10.1111/cogs.12413

Lehet, M., & Holt, L. L. (2020). Nevertheless, it persists: Dimension-based statistical learning and normalization of speech impact different levels of perceptual processing. Cognition, 202, 104328. https://doi.org/10.1016/j.cognition.2020.104328

Liu, R., & Holt, L. L. (2015). Dimension-based statistical learning of vowels. Journal of Experimental Psychology. Human Perception and Performance, 41(6), 1783–1798. https://doi.org/10.1037/xhp0000092

Maiste, A. C., Wiens, A. S., Hunt, M. J., Scherg, M., & Picton, T. W. (1995). Event-Related Potentials and the Categorical Perception of Speech Sounds. Ear and Hearing, 16(1), 68–89. https://doi.org/10.1097/00003446-199502000-00006

Makeig, S., Müller, M. M., & Rockstroh, B. (1996). Effects of voluntary movements on early auditory brain responses. Experimental Brain Research, 110(3), 487–492. https://doi.org/10.1007/bf00229149

Maris, E., & Oostenveld, R. (2007). Nonparametric statistical testing of EEG-and MEG-data. Journal of Neuroscience Methods, 164(1), 177–190.

Maye, J., Weiss, D. J., & Aslin, R. N. (2008). Statistical phonetic learning in infants: facilitation and feature generalization. Developmental Science, 11(1), 122–134. https://doi.org/10.1111/j.1467-7687.2007.00653.x

Mesgarani, N., Cheung, C., Johnson, K., & Chang, E. F. (2014). Phonetic feature encoding in human superior temporal gyrus. Science (New York, N.Y.), 343(6174), 1006–1010. https://doi.org/10.1126/science.1245994

Moberly, A. C., Bhat, J., Welling, D. B., & Shahin, A. J. (2014). Neurophysiology of spectrotemporal cue organization of spoken language in auditory memory. Brain and Language, 130, 42–49. https://doi.org/10.1016/j.bandl.2014.01.007

Mulert, C., Pogarell, O., Juckel, G., Rujescu, D., Giegling, I., Rupp, D., Mavrogiorgou, P., Bussfeld, P., Gallinat, J., Möller, H. J., & Hegerl, U. (2004). The neural basis of the P300 potential. European Archives of Psychiatry and Clinical Neuroscience, 254(3), 190–198. https://doi.org/10.1007/s00406-004-0469-2

Näätänen, R. (2008). Mismatch negativity (MMN) as an index of central auditory system plasticity. International Journal of Audiology, 47(sup2), S16–S20. https://doi.org/10.1080/14992020802340116

Näätänen, R., & Alho, K. (1997). Mismatch Negativity – The Measure for Central Sound Representation Accuracy. Audiology and Neurotology, 2(5), 341–353. https://doi.org/10.1159/000259255

Näätänen, R., Paavilainen, P., Rinne, T., & Alho, K. (2007a). The mismatch negativity (MMN) in basic research of central auditory processing: A review. Clinical Neurophysiology, 118(12), 2544–2590. https://doi.org/10.1016/j.clinph.2007.04.026

Näätänen, R., Paavilainen, P., Rinne, T., & Alho, K. (2007b). The mismatch negativity (MMN) in basic research of central auditory processing: a review.

Näätänen, R., & Picton, T. (1987). The N1 wave of the human electric and magnetic response to sound: a review and an analysis of the component structure. Psychophysiology, 24(4), 375–425. https://doi.org/10.1111/j.1469-8986.1987.tb00311.x

Nichols, T. E., & Holmes, A. P. (2002). Nonparametric permutation tests for functional neuroimaging: A primer with examples. Human Brain Mapping, 15(1), 1–25. https://doi.org/10.1002/hbm.1058

Nieuwenhuis, S., Aston-Jones, G., & Cohen, J. D. (2005). Decision Making, the P3, and the Locus Coeruleus-Norepinephrine System. Psychological Bulletin, 131(4), 510–532. https://doi.org/10.1037/0033-2909.131.4.510

Norris, D., McQueen, J., & Cutler, A. (2000). Merging information in speech recognition: feedback is never necessary. The Behavioral and Brain Sciences, 23(3), 299–325; discussion 325-70. https://doi.org/10.1017/s0140525x00003241

Pantev, C., Hoke, M., Lehnertz, K., Lütkenhöner, B., Fahrendorf, G., & Stöber, U. (1990). Identification of sources of brain neuronal activity with high spatiotemporal resolution through combination of neuromagnetic source localization (NMSL) and magnetic resonance imaging (MRI). Electroencephalography and Clinical Neurophysiology, 75(3), 173–184.

Pereira, O., Gao, Y. A., & Toscano, J. C. (2018). Perceptual Encoding of Natural Speech Sounds Revealed by the N1 Event-Related Potential Response. Auditory Perception & Cognition, 1(1–2), 112–130. https://doi.org/10.1080/25742442.2018.1545106

Samuel, A. G., & Kraljic, T. (2009). Perceptual learning for speech. Attention, Perception & Psychophysics, 71(6), 1207–1218. https://doi.org/10.3758/app.71.6.1207

Scherg, M., & Von Cramon, D. (1985). Two bilateral sources of the late AEP as identified by a spatio-temporal dipole model. Electroencephalography and Clinical Neurophysiology/Evoked Potentials Section, 62(1), 32–44.

Schertz, J., Cho, T., Lotto, A., & Warner, N. (2016). Individual differences in perceptual adaptability of foreign sound categories. Attention, Perception & Psychophysics, 78(1), 355–367. https://doi.org/10.3758/s13414-015-0987-1

Schertz, J., & Clare, E. J. (2019). Phonetic cue weighting in perception and production. WIREs Cognitive Science, 11(2), e1521. https://doi.org/10.1002/wcs.1521

Schwab, E., Nusbaum, H., & Pisoni, D. (1985). Some effects of training on the perception of synthetic speech. Human Factors, 27(4), 395–408. https://doi.org/10.1177/001872088502700404

Tang, C., Hamilton, L. S., & Chang, E. F. (2017). Intonational speech prosody encoding in the human auditory cortex. Science, 357(6353), 797–801. https://doi.org/10.1126/science.aam8577

Toscano, J. C., Anderson, N. D., Fabiani, M., Gratton, G., & Garnsey, S. M. (2018). The time-course of cortical responses to speech revealed by fast optical imaging. Brain and Language, 184, 32–42. https://doi.org/10.1016/j.bandl.2018.06.006

Toscano, J. C., McMurray, B., Dennhardt, J., & Luck, S. J. (2010). Continuous perception and graded categorization: electrophysiological evidence for a linear relationship between the acoustic signal and perceptual encoding of speech. Psychological Science, 21(10), 1532–1540. https://doi.org/10.1177/0956797610384142

Viswanathan, V., Bharadwaj, H. M., Shinn-Cunningham, B. G., & Heinz, M. G. (2021). Modulation masking and fine structure shape neural envelope coding to predict speech intelligibility across diverse listening conditions. The Journal of the Acoustical Society of America, 150(3), 2230–2244. https://doi.org/10.1121/10.0006385

Vroomen, J., Linden, S. van, Gelder, B. de, & Bertelson, P. (2007). Visual recalibration and selective adaptation in auditory-visual speech perception: Contrasting build-up courses. Neuropsychologia, 45(3), 572–577. https://doi.org/10.1016/j.neuropsychologia.2006.01.031

Winn, M. B., Chatterjee, M., & Idsardi, W. J. (2013). Roles of Voice Onset Time and F0 in Stop Consonant Voicing Perception: Effects of Masking Noise and Low-Pass Filtering. Journal of Speech, Language, and Hearing Research, 56(4), 1097–1107. https://doi.org/10.1044/1092-4388(2012/12-0086)

Wu, Y. C., & Holt, L. L. (2022). Phonetic Category Activation Predicts the Direction and Magnitude of Perceptual Adaptation to Accented Speech. Journal of Experimental Psychology: Human Perception and Performance, 48(9), 913–925. https://doi.org/10.1037/xhp0001037

Yingling, C. D., & Hosobuchi, Y. (1984). A subcortical correlate of P300 in man. Electroencephalography and Clinical Neurophysiology/Evoked Potentials Section, 59(1), 72–76. https://doi.org/10.1016/0168-5597(84)90022-4

Zhang, H., Wiener, S., & Holt, L. L. (2022). Adjustment of cue weighting in speech by speakers and listeners: Evidence from amplitude and duration modifications of Mandarin Chinese tone. The Journal of the Acoustical Society of America, 151(2), 992–1005. https://doi.org/10.1121/10.0009378

Zhang, X., & Holt, L. L. (2018). Simultaneous tracking of coevolving distributional regularities in speech. Journal of Experimental Psychology: Human Perception and Performance, 44(11), 1760–1779. https://doi.org/10.1037/xhp0000569

Zhang, X., Wu, Y. C., & Holt, L. L. (2021). The Learning Signal in Perceptual Tuning of Speech: Bottom Up Versus Top-Down Information. Cognitive Science, 45(3), e12947. https://doi.org/10.1111/cogs.12947

